# Comparative analysis of polymerase chain reaction (PCR) assays in detection of *Orientia tsutsugamushi* from mite vectors

**DOI:** 10.64898/2026.09.28.754975

**Authors:** R. Govindarajan, S. Gowri Sankar, R. Manju, P. Philip Samuel, A. Alwin Prem Anand

**Author notes:** Corresponding authors P. Philip Samuel, Unit of Vector-borne and Zoonotic diseases, ICMR-NIVCR Filed station, Madurai - 625002, India, A. Alwin Prem Anand, Department of Biotechnology - Bioinformatics Infrastructure Facilities (DBT-BIF) Centre, (Under DBT Biotechnology Information System Network (BTISNet) Scheme), Lady Doak College, Madurai - 625002, India. Equally contributed (R. Govindarajan, S. Gowri Sankar).

## Abstract

*Orientia tsutsugamushi* is the causative agent of scrub typhus infection in humans. This pathogen is transmitted into mammalian hosts including humans through *Leptotrombidium* mites. Till date, detection of *O. tsutsugamushi* in human was performed using PCR assay with 56kDa type specific antigen (TSA56), 47kDa outer membrane protein (47kDa-OMP) and groEL chaperonin gene. To understand the efficiency of different genes for the detection of *O. tsutsugamushi* in *Leptotrombidium* mites, we performed a comparative PCR assay with various universal primers targeting different genes including TSA56, 47kDa-OMP and groEL gene. We have collected 500 *Leptotrombidium* mites from Vellore, Tamil Nadu and, the mites were segregated into 50 pools with 10 mites in each pool were used for DNA extraction. The molecular detection of *O. tsutsugamushi* was performed using PCR assays (including conventional, nested and duplex PCR). The comparative PCR analysis showed nested PCR of groEL gene has high efficiency in detecting the presence of *O. tsutsugamushi* in chigger samples compared to other genes. Interestingly, groEL target demonstrated high efficiency for detection of *O. tsutsugamushi* in chigger samples and, when used with nested primers, can facilitate differentiation of *O. tsutsugamushi* from other rickettsial pathogens. Thus, we conclude groEL gene nested PCR assay can be used routinely to detect *O. tsutsugamushi* in trombiculid mites.

## Introduction

Scrub typhus is a form of typhus caused by the intracellular parasite *Orientia tsutsugamushi* which is a gram-negative α-proteobacterium comes under Rickettsiaceae family. The bacterium was first isolated and identified in Japan in 1930 (Nagayo et al. 1930). Based on review article by Bonell and colleagues (Bonell et al. 2017), seroprevalence ranged from 9.3 to 27.9% with a median of 22.2% among six countries across Asia and, 6% of Scrub typhus will die if untreated, while 1.5% if treated and mortality can reach up to 13% where treatment does not work. Complication including brain infection (Dittrich et al. 2015) and multiple organ failure (Griffith et al. 2014) can increase the mortality rate to 14% and 24% respectively. Pregnancy related complication has also been reported in Scrub typhus infection (Chansamouth et al. 2016; McGready et al. 2010; McGready et al. 2014).

The disease is transmitted to humans by mites of Trombiculidae family and humans are accidental hosts in this zoonosis (Watt and Parola 2003). The transmission of scrub typhus in the tropical areas occurs throughout the year; while in temperate zones, it is seasonal. So far in *Trombiculidae, O. tsutsugamushi* was detected from 46 species of mite species (Elliott et al. 2019). Till date, laboratory diagnosis of *O. tsutsugamushi* is done using various techniques - IFA, ELISA, PCR (including nested, conventional, multiplex and real-time). Though IFA is considered as a gold standard; the requirement of high cost and sophisticated instrument makes it a tougher to use in regular laboratory setup as well as there was no uniform positivity cutoff titre reported across countries (Blacksell et al. 2007; Saraswati et al. 2019). Owing to this, inexpensive PCR methods based on a variety of genes like 56-kDa, 47-kDa and GroEL are used (Janardhanan et al. 2014; Kim et al. 2011; Paris et al. 2009; Park et al. 2005). As most researchers assesses the usefulness of these genes in detecting *O. tsutsugamushi* from human samples (Elliott et al. 2021; Fournier et al. 2008; Kim et al. 2011; Kim et al. 2006), few research was carried on in assessing the applicability of these genes in detecting the pathogen from the zoonotic carriers. Hence, the study was aimed to find out the applicability of these widely used genes in detecting the pathogen from its carriers as it is necessary for epidemiological purposes as well as for identifying potential hotspots for the spread of the disease.

## Materials and Methods

### Chigger mite collection and identification

Chigger mites collected from small mammals (rodents and shrews) in Vellore, Tamil Nadu, India, during 2017–2018 were preserved in 80% ethanol until further processing. A total of 500 individual chiggers were examined for species-level identification based on diagnostic morphological characteristics of the gnathosoma, legs, idiosoma, and scutum.

Morphological examination was performed using an autofluorescence-based approach adapted from Kumlert et al. (2018). Individual chiggers were temporarily positioned between coverslips in sterile water without the addition of fluorescent dyes and examined using a Magnus MLXi+ microscope equipped with an epifluorescence illuminator and fluorescein isothiocyanate (FITC) filter, together with bright-field illumination. Species identification was based on the characteristic differential autofluorescence patterns of the scutum and other diagnostically relevant integumentary structures, assessed in conjunction with bright-field observations. Morphometric measurements of the scutum were documented using MagVision imaging software following the procedures described by Kumlert et al. (2018).

Following morphological examination, identified specimens were recovered from the temporary mounts and pooled according to species and retained in ethanol for subsequent molecular analysis. No permanent mounting medium or chemical clearing procedure was applied, thereby permitting the specimens to be used for DNA extraction and PCR.

### DNA extraction and PCR

Mite pools were homogenized using electric homogenizer and DNA was extracted using DNeasy Blood & Tissue Kit (Qiagen), with slight modifications. After homogenization with PBS pH 7.2 (50 mM potassium phosphate, 150 mM NaCl), 20 μl proteinase K and 200 μl Buffer AL was added and incubated at 56°C for 4 hours. Further extraction procedure was followed by established protocol. PCR was performed in thirty µl of reaction mixture contained 15 µl Taq 2x Master Mix red containing 1.5 mM MgCl_2_ (Ampliqon), 1 µl each of forward and reverse primers, 5 µl of DNA template. All the PCR products were resolved in agarose gel electrophoresis with appropriate molecular weight markers. PCR primers, targeted genes, type of PCR and annealing conditions are given in Table 1.

**Table 1.**
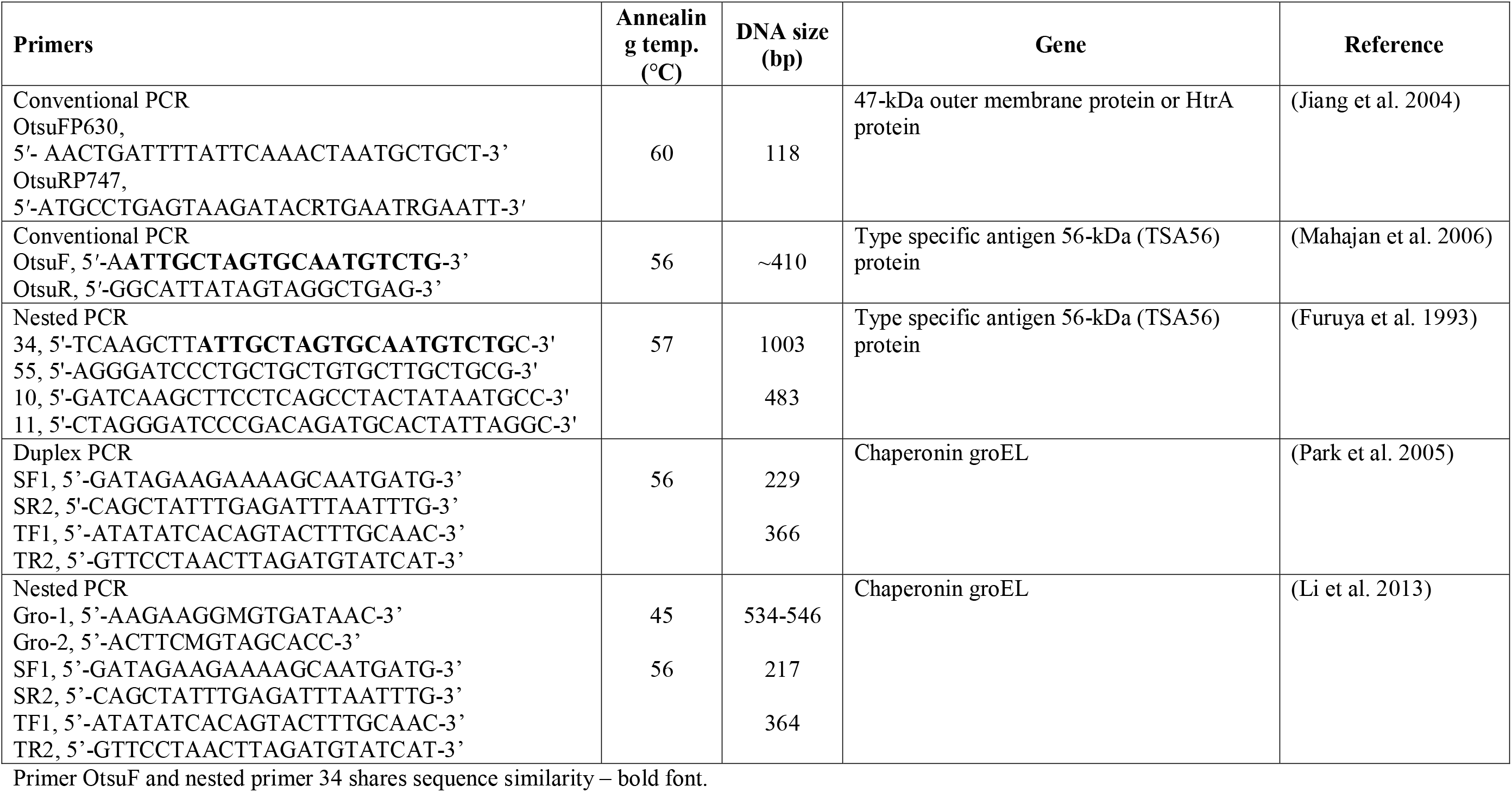
Genes and primers used in this study

## Result

The majority of the pooled samples belongs to *Leptotrombidium deliense* (42/50 pools), followed by *L. indicum* (4/50), *L. rajasthanense* (3/50) and *L. insigne* (1/50). Among these trombiculid chigger mites, *L. deliense* and *L. rajasthanense* were positive for the presence of scrub typhus pathogen, *O. tsutsugamushi* (Table 2). Interestingly, the *O. tsutsugamushi* positivity in these chigger mite same pooled samples vary with the primers used (Fig. 1, Table 3). The amplification of groEL genes using nested PCR primers shows positivity in 14 pools (*L. deliense* –13/14, *L. rajasthanense* – 1/14); while, duplex PCR primers show positivity only in 9 pools (*L. deliense* – 8/9, *L. rajasthanense* – 1/9). Amplification of 56-kDa gene using nested PCR (primers 34, 55, 10 & 11) shows positivity in 8 pools (*L. deliense* – 7/8, *L. rajasthanense* – 1/8); while conventional PCR (OtsuF/OtsuR primer) shows only positivity in 4 pools of *L. deliense* samples. The OtsuFP630/OtsuRP747 amplifying 47-kDa gene shows positivity in 5 pools (*L. deliense* – 4/5, *L. rajasthanense* – 1/5) (Fig. 1, Table 2&3). None of chiggers are positive for *Rickettsia* species.

**Table 2.** Presence of *O. tsutsugamushi* tested in ectoparasites using different genes and PCR

| Trombiculid mite | Number of Pools | Number of pools positive for *O. tsutsugamushi* |  |  |  |  |
| --- | --- | --- | --- | --- | --- | --- |
|  |  | 47-kDa gene | groEL gene |  | Type specific antigen 56-kDa (TSA56) gene |  |
|  |  | Conventional PCR – OtsuFP630/OtsuRP747 | Nested PCR – Gro1/Gro2, TF1/TR2 & SF1/SR2 | Duplex PCR – SF1/SR2 & TF1/TR2 | Conventional PCR – OtsuF/OtsuR | Nested PCR – 34/55 & 10/11 |
| *Leptotrombidium insigne* | 1 | 0 | 0 | 0 | 0 | 0 |
| *Leptotrombidium deliense* | 42 | 4 | 13 | 8 | 4 | 7 |
| *Leptotrombidium indicum* | 4 | 0 | 0 | 0 | 0 | 0 |
| *Leptotrombidium rajasthanense* | 3 | 1 | 1 | 1 | 0 | 1 |
| **Grand Total** | **50** | **5** | **14** | **9** | **4** | **8** |

**Table 3:** Positivity of target genes in different combination

| Trombiculid mite | Number of positive pools |  |  |  |  |  |
| --- | --- | --- | --- | --- | --- | --- |
|  | All PCR positive | Conv. 47-kDa + Nested 56-kDa + Duplex groEL + Nested groEL | Nested 56-kDa + Duplex groEL + Nested groEL | Conv. 56-kDa + Duplex groEL + Nested groEL | Nested groEL alone | Total |
| *Leptotrombidium deliense* | 3 | 1 | 3 | 1 | 5 | 13 |
| *Leptotrombidium rajasthanense* | 0 | 1 | 0 | 0 | 0 | 1 |
| **Grand total** | **3** | **2** | **3** | **1** | **5** | **14** |

**Fig 1:**
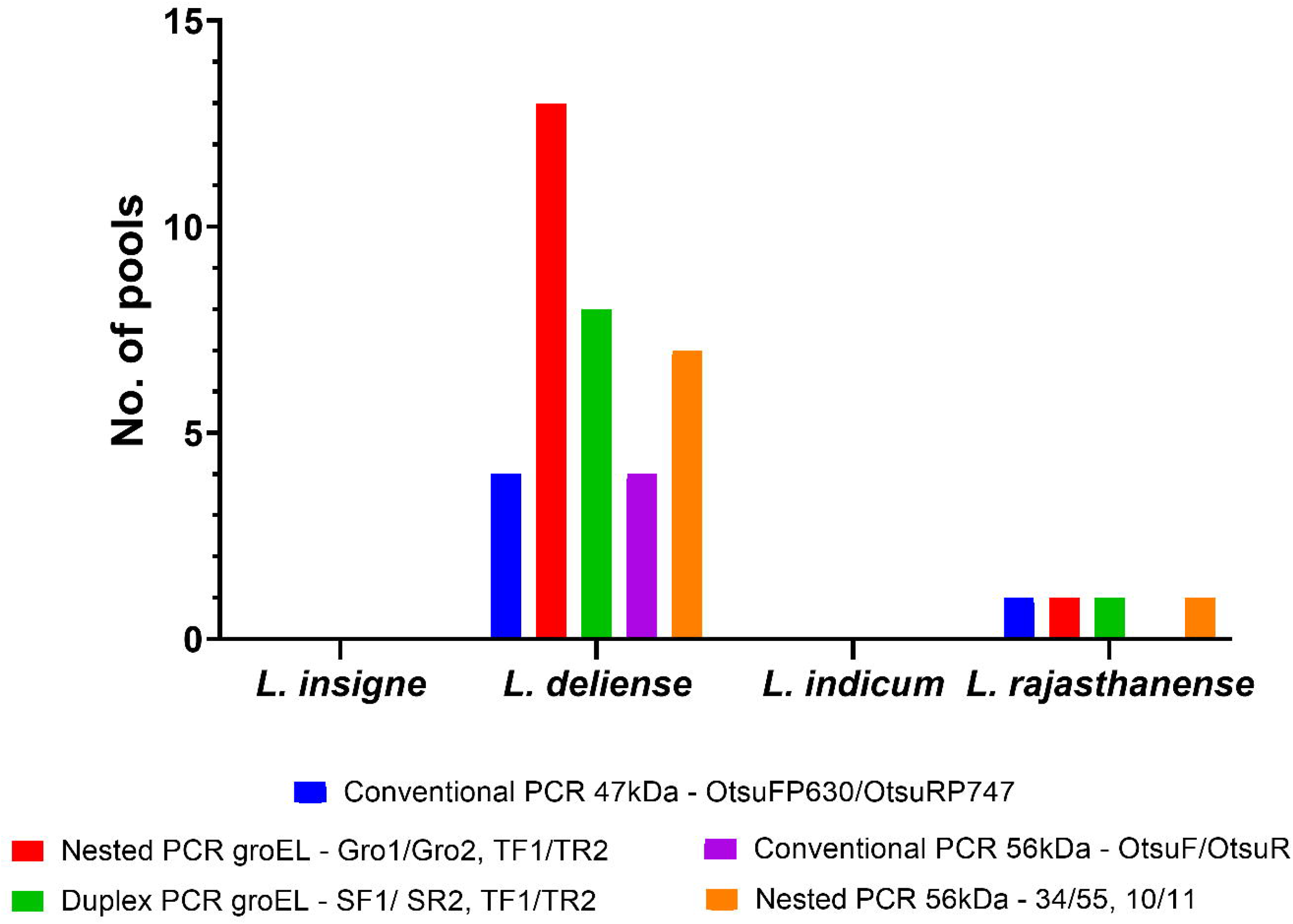
Positivity for *O. tsutsugamushi* in *Leptotrombidium* species using different genes and PCR amplification

## Discussion

*O. tsutsugamushi* is transmitted by chigger mites belonging to the family Trombiculidae, with the majority of known vector species belonging to the genus *Leptotrombidium* (Elliott et al. 2019). Among the *Leptotrombidium* species, *L. akamushi, L. arvinum, L. bodense, L. deliense, L. fletcheri, L. fuji, L. intermedium, L. keukenshrijveri, L. pallidum, L. palpale, L. pavlovskyi, L. peniculatum, L. scutellare, L. vivericola*, (Elliott et al. 2019; Mullen and Oconnor 2019), *L. chaigraiensis, L. dihumerale, L. gaohuensis, L. himizu, L. insulare* and *L. zetum* were reported to be positive for *O. tsutsugamushi* (Moniuszko et al. 2022).

In India, 204 species of trombiculid chigger mites have been reported, of which 55 belong to the genus *Leptotrombidium* (Fernandes and Kulkarni 2003). Among these, 16 species namely *L. bhattipadense, L. dehradunense, L. deliense, L. delimushi, L. discrepans, L. fulmentum, L. imphalum, L. indicum, L. insigne, L. jayewickremei* (currently listed under genus *Ericotrombidium* (Womersley, 1952) (Moniuszko et al. 2022)), *L. keukenschrijveri, L. kulkarnii, L. spilletti, L. pseudogliricolens* and *L. rajasthanense*, have been reported from Tamil Nadu (Candasamy et al. 2016; Paulraj et al. 2022; Renu et al. 2021). More recently, two additional species *L. puta* and *L. langati*, have also been reported from Tamil Nadu (Renu et al. 2026). *L. deliense* is considered as the major vector in India and Southeast Asia (Elliott et al. 2019). In India, *O. tsutsugamushi* has also been detected in *L. imphalum* (Devamani et al. 2026), *L. rajasthanense* and *L. jayawickremei* (Prakash et al. 2022) suggesting that the association between *O. tsutsugamushi* and chigger mites extends beyond *L. deliense*.

The present study builds upon the chigger mite specimens collected from small mammals (rodents and shrews) in Vellore, Tamil Nadu, India, during 2017–2018. These specimens were previously investigated by Paulraj et al. (2022), in which chigger mites were identified morphologically using microscopic examination and standard taxonomic keys. In the present study, a separate set of specimens from the same collection was examined using the morphological criteria described by Kumlert et al. (Kumlert et al. 2018), followed by PCR amplification for the detection and molecular identification of *O. tsutsugamushi*. Thus, the present study extends the earlier morphological investigation by incorporating a molecular approach to assess the association of *O. tsutsugamushi* with chigger mites collected from the study area.

In the present study, we report the presence of *L. deliense, L. indicum, L. rajasthanense* and *L. insigne* chiggers in small mammals from Vellore, which is also been reported by Paulraj et al., (2022) in the same study area. Among these species, *L. deliense* and *L. rajasthanense* chiggers were positive for *O. tsutsugamushi. L. deliense* was frequently positive for *O. tsutsugamushi* across South Asia including India, Southeast Asia, East Asia and Oceania/Pacific (Elliott et al. 2019; Moniuszko et al. 2022). Notably, *L. rajasthanense* was reported to be positive for *O. tsutsugamushi* in Vellore based on amplification of the *traD* gene (Prakash et al. 2022), whereas in the present study, *L. rajasthanense* was positive based on amplification of the *groEL* gene. The detection of *O. tsutsugamushi* in *L. rajasthanense* using a different molecular target therefore provides additional molecular evidence of its association with *O. tsutsugamushi* in Vellore.

### 47-kDa (Omp or HtrA) gene

The 47-kDa protein of *O. tsutsugamushi* is a conserved outer membrane protein (Omp) contains a trypsin domain and has sequence homology to human HtrA (high temperature requirement A) serine protease protein family (Chen et al. 2011). In our observation, amplification of 47-kDa gene using OtsuFP630/OtsuRP747 primer shows least positivity (5/70 pools) for *O. tsutsugamushi* presence in the ectoparasites. According to Jiang and colleagues (Jiang et al. 2013), though 47-kDa gene is conserved there was a 3.3% divergence within other strains with exception of *O. chuto*. The divergence of 47-kDa gene of *O. chuto* strain ranged from 17.7-18.2% with other strains of *O. tsutsugamushi* (Izzard et al. 2010). The reverse primer (OtsuRP747) of 47-kDa gene was designed as degenerate primer based on the sequences of the *O. tsutsugamushi* Karp (L31934), Kato (L11697), Gilliam (L31933), Boryong (L319335)(Jiang et al. 2004), which helps in amplifying possible combination of nucleotides for a coding sequence.

We performed an *in silico* primer-binding analysis of the 56-kDa gene using 18 available reference genome sequences (Fig. 2; Table 4). Interestingly, *in silico* analysis of the conventional OtsuFP630/OtsuRP747 primer pair showed 100% sequence identity at the OtsuFP630 binding site across all 18 genomes. In contrast, the OtsuRP747 primer showed an exact match without mismatches only in a few strains (K4-135, CH219, Kato, and Ikeda), whereas the other isolates exhibited varying numbers of mismatches at the reverse-primer binding site (Fig. 2). These sequence variations may contribute to reduced amplification efficiency and, consequently, lower detection rates of *O. tsutsugamushi*.

**Table 4:** Details on sequences used for BLAST and alignment

| Sequence ID | Organism | Strain name | Source | Collection place | Year isolated |
| --- | --- | --- | --- | --- | --- |
| NZ_CM176317.1 | *O. tsutsugamushi* | K4-135 | Human blood | South Korea: Gyeonggi-do | 2024 |
| NZ_CM176316.1 | *O. tsutsugamushi* | CH219 | Human blood | South Korea: Kangwon-do | 2023 |
| NZ_CP044031.1 | *O. tsutsugamushi* | Wuj/2014 | Blood from febrile patient | China:Zhejiang province | 2014 |
| NZ_LS398547.1 | *O. tsutsugamushi* | UT176 | Blood from patient | Thailand* | 2004 |
| NZ_LS398552.1 | *O. tsutsugamushi* | UT76 | Blood from patient | Thailand* | 2003 |
| NZ_CP142420.1 | *O. tsutsugamushi* | TW-22 | Blood from patient | Taiwan: Kaohsiung City | 2007 |
| NZ_CP142421.1 | *O. tsutsugamushi* | TW-1 | Blood from patient | Taiwan: Lienchiang County | 2007 |
| NZ_LS398549.1 | *O. tsutsugamushi* | TA686 | *Tupaia glis* * | Thailand* | 1963* |
| NZ_LS398550.1 | *O. tsutsugamushi* | Kato | Blood from patient* | Japan* | 1955* |
| NZ_LS398548.1 | *O. tsutsugamushi* | Karp | Blood from patient* | New Guinea* | 1943* |
| NZ_CP166958.1 | *O. tsutsugamushi* | JJOtsu8 | Human PBMC | India: Vellore | 2023 |
| NZ_CP166957.1 | *O. tsutsugamushi* | JJOtsu7 | Human PBMC | India: Vellore | 2024 |
| NZ_CP166956.1 | *O. tsutsugamushi* | JJOtsu6 | Human PBMC | India: Vellore | 2023 |
| NZ_CP166955.1 | *O. tsutsugamushi* | JJOtsu5 | Human PBMC | India: Vellore | 2023 |
| NZ_CP166954.1 | *O. tsutsugamushi* | JJOtsu1 | Human PBMC | India: Vellore | 2022 |
| NZ_LS398551.1 | *O. tsutsugamushi* | Gilliam | Blood from patient* | Burma* | 1943* |
| NC_010793.1 | *O. tsutsugamushi* | Ikeda | Blood from patient† | Japan: Niigata† | 1979† |
| NC_009488.1 | *O. tsutsugamushi* | Boryong | Blood from patient* | South Korea: Seoul | Late 1980s* |
| NC_010263.3 | *R. rickettsii* | Lowa | - | - | - |

\*Details taken from Minahan et al., (2024) †Details taken from Nakayama et al., (2008)

**Fig 2:**
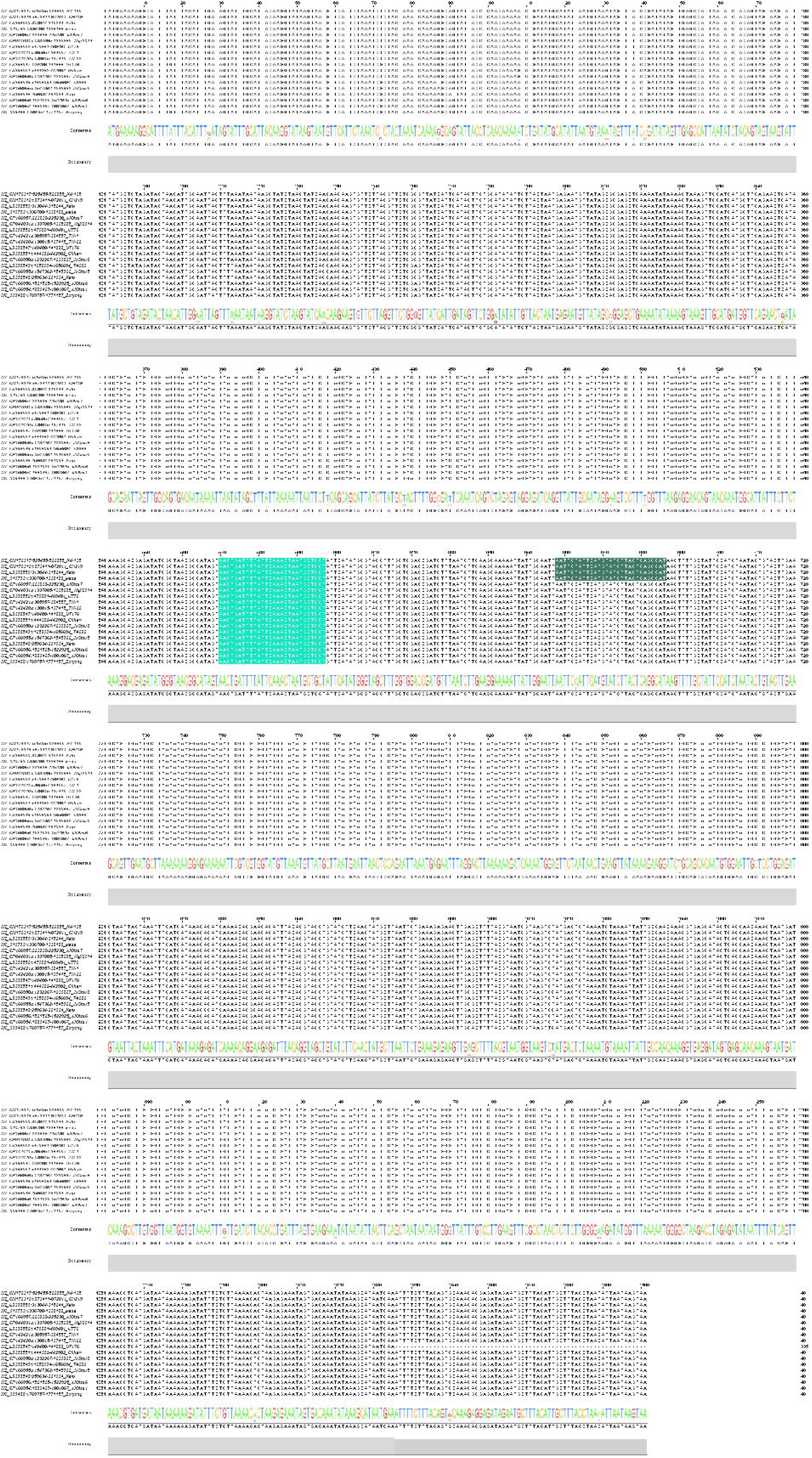
PCR primer-binding sites on the 47-kDa gene across 18 *O. tsutsugamushi* genome sequences. Strain information is provided in Table 4. The binding sites of the conventional primers OtsuFP630 and OtsuRP747 are indicated in different shades of green.

In our study, the lower positivity observed with the 47-kDa gene may be attributable to one or both of the following factors: (1) extensive sequence variation or mutations within the primer-binding region that exceed the binding capacity of the degenerate primer, and/or (2) a low copy number of the target gene in the samples.

### 56-kDa (OmpA or TSA56) gene

The 56-kDa is a outer-membrane protein of *O. tsutsugamushi*, which is highly immunogenic and recognized by antibodies in most scrub-typhus patients (Chen et al. 2011). Upon analysing for the presence of *O. tsutsugamushi* in ectoparasites using 56-kDa gene, we have observed conventional PCR (OtsuF/OtsuR) showing less positivity than nested PCR (34, 55, 11 & 10) within the same pools of ectoparasite samples. Reports shows positive rate of OtsuF/OtsuR primers about 14.2% (3/21) (Mahajan et al. 2006), 17.5% (11/63) (Parola et al. 2008) and 95.8% (23/24) (Janardhanan et al. 2014). Notably, OtsuF/OtsuR primers in comparison with 16S rRNA primers showed less positivity (Kim et al. 2016; Sonthayanon et al. 2006). Janardhanan and colleagues (Janardhanan et al. 2014) have performed a comparative study using the same primers and, reported that 95.8% (23/24 samples) positivity in conventional primers compared to nested primers with 75% (18/24 samples) positivity rate; which is contrast to our observation. Kim and colleagues (Kim et al. 2011) used primers 10/11 for conventional PCR and 34, 55, 10 & 11 for nested PCR and found that conventional PCR was negative for 41 scrub typhus patient samples, while nested PCR was 87.8% positive in the same samples; which is similar to our observation, where conventional PCR shows less positivity than nested PCR within the same pools.

We performed an *in silico* primer-binding analysis of the 56-kDa gene using 18 available reference genome sequences (Fig. 3, Table 4). We observed 100% consensus sequence identity for the OtsuF and primer 34 binding sites in all strains based on BLAST analysis. However, nucleotide mismatches were observed at the OtsuR and primer 10 binding sites in some strains (Boryong, JJOtsu8, and TA686). Furthermore, sequence variations at the reverse primer 11 and 55 binding sites may lead to reduced identification of *O. tsutsugamushi* (Fig. 3). It is possible that these sequence variations may interfere with primer binding and amplification, resulting in an insufficient number of amplicon copies for visualization on an agarose gel. This may explain the failure of PCR with the primer 10/11 combination and the lower positivity observed in the nested PCR reported by Janardhanan and colleagues.

**Fig 3:**
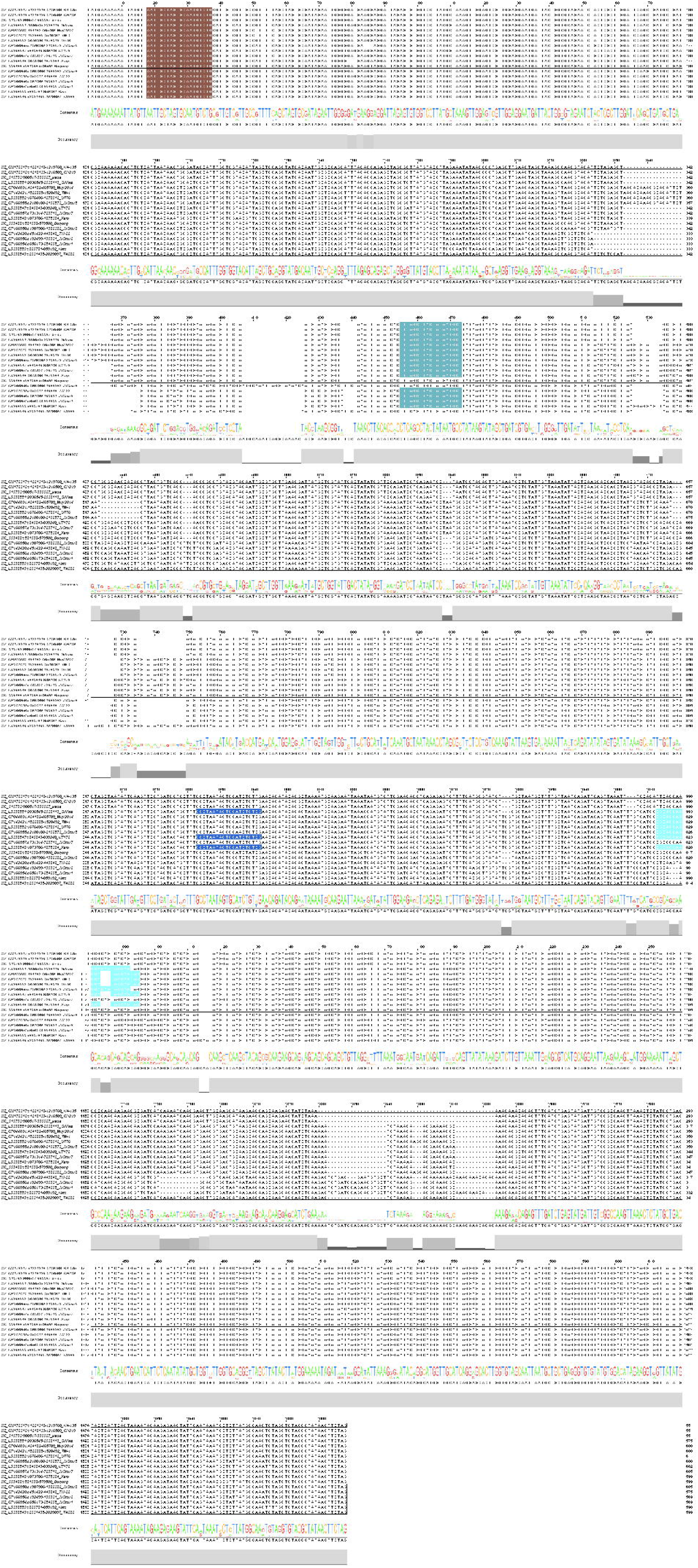
PCR primer-binding sites on the 56-kDa gene across 18 *O. tsutsugamushi* genome sequences. Strain information is provided in Table 4. Brown indicates the OtsuF and primer 34 binding sites; light blue indicates the OtsuR and primer 10 binding sites; dark blue indicates the primer 11 binding site; and pale blue indicates the primer 55 binding site.

Additionally, the genome sequences of the Indian strains (JJOtsu1, JJOtsu5, JJOtsu6, JJOtsu7, and JJOtsu8) showed mismatches within the reverse-primer binding sites. These mismatches may interfere with amplification of the target gene and consequently result in failure to detect *O. tsutsugamushi* in infected hosts. Among the Indian strains analysed, only JJOtsu5 was predicted to be amplifiable using the nested primers 34 and 55 (Fig. 3). Taken together, this may explain the lower positivity observed in our study compared with that obtained using the other genes evaluated.

### groEL gene

The groEL is a heat-shock protein (HSP60/chaperonin 60), which have higher divergence among the family Rickettsiaceae and can differentiate the genus *Rickettsia* and *Orientia* (Lee et al. 2003; Park et al. 2005). The higher positivity rate observed for the *groEL* gene compared with other target genes (47-kDa and 56-kDa), highlights the suitability of this target for detecting *O. tsutsugamushi* in ectoparasite samples. The superior performance of *groEL* nested PCR compared with duplex PCR (Table 2) is likely attributable to the high conservation of the *groEL* gene among *O. tsutsugamushi* strains and its greater divergence from *Rickettsia* species, as reported previously (Lee et al. 2003; Li et al. 2013; Park et al. 2005). These findings are consistent with earlier studies demonstrating higher conservation of *groEL* (Arai et al. 2013; Paris et al. 2009) among *O. tsutsugamushi* strains.

In studies using duplex PCR, *O. tsutsugamushi* was reported in 1.34% (7/519) *Rattus norvegicus* samples (Hotta et al. 2016) and in 1.4% (2/141 pools) chigger mite samples (Binh et al. 2020). In a clinical study, 47 of 52 (90%) patients were found positive for *O. tsutsugamushi* using nested PCR for groEL which was also positive for IFA against *O. tsutsugamushi* antigen (Li et al. 2013). Similarly, a PCR-based study of 140 scrub typhus IgM-positive cases reported a positivity rate of 61.42% (Anitharaj et al. 2020). In a previous study from Madurai, *O. tsutsugamushi* was detected by nested PCR targeting the *groEL* gene in blood samples of small mammals, including *R. rattus, Suncus murinus* and *Bandicota bengalensis*, as well as in several non-trombiculid ectoparasites, including *Oribatida* sp., *Dermanyssus gallinae, Xenopsylla astia, X. cheopis, Ctenophthalis* sp., *C. felis, Rhipicephalus sanguineus* and *R. haemaphysaloides* (Govindarajan et al. 2024). Notably, all chigger mites examined in that study were negative for *O. tsutsugamushi*. In contrast, *O. tsutsugamushi* was detected in *L. deliense* and *L. rajasthanense* from Vellore using a multicopy *traD*-targeting assay (Prakash et al. 2022) and in this study (Vellore), supporting the potential involvement in the transmission cycle of scrub typhus. In this study, nested PCR shows higher positivity, it might be due to the first-round Gro1/Gro2 amplification provides a broad, conserved *groEL* target, followed by species-specific second-round amplification with TF1/TF2 for *O. tsutsugamushi*.

### Comparative analysis using different genes and PCR

There were very few comparative studies using different genes and PCR for the diagnosis of *O. tsutsugamushi*. Candasamy and colleagues (2016) performed a study using 56-kDa primer (same as in this study), and a different primer for groEL, they observed 4% (2/50) rat samples were positive using groEL gene, while 56-kDa does not yield any results. Another study from the same group (Sadanandane et al. 2021) using same primers as in this study, showing in rodent blood samples 1.49% and 15.47% positive for nested PCR 56-kDa gene and groEL gene, respectively. In a clinical study, Anitharaj and colleagues (2020) used nested PCR for 56-kDa, 47-kDa and groEL genes and, among 140 scrub typhus IgM ELISA positive cases 30%, 51.42% and 61.42% were positive for 56-kDa, 47-kDa and groEL, respectively.

In the present study, duplex and nested PCR for groEL gene has shown high positivity range alone or in combination of other genes (Table 2 & 3) may be attributed to the design of the *groEL*-targeting assay and the additional amplification step. *In silico* analysis of the groEL primer-binding sites across 18 different strains of *O. tsutsugamushi* and a *Rickettsia rickettsii* strain (Table 4) using duplex and nested PCR primers showed that: (1) the degenerate primers Gro-1 and Gro-2 can be used for the identification of *Rickettsia* and *Orientia* species; (2) the nested primers SF1 and SF2 bind only to *Rickettsia* species; and (3) the nested primers TF1 and TF2 bind only to *O. tsutsugamushi* (Fig. 4). Importantly, *in silico* analysis of the primer-binding regions across 18 *O. tsutsugamushi* strains showed no sequence variation or mutations at the primer-binding sites, indicating high conservation of the target regions. Thus, the conserved nature of the Gro1/Gro2 target sites, followed by species-specific amplification with TF1/TF2, may enhance the ability of the nested PCR assay to detect low levels of *O. tsutsugamushi* DNA in samples. (Fig. 4). The increased positivity observed in the present study may therefore reflect, at least in part, the broad initial amplification and subsequent species-specific enrichment inherent to the nested-PCR approach. Taken together, these findings suggest that the groEL gene is a useful target for the detection and identification of *O. tsutsugamushi*.

**Fig 4:**
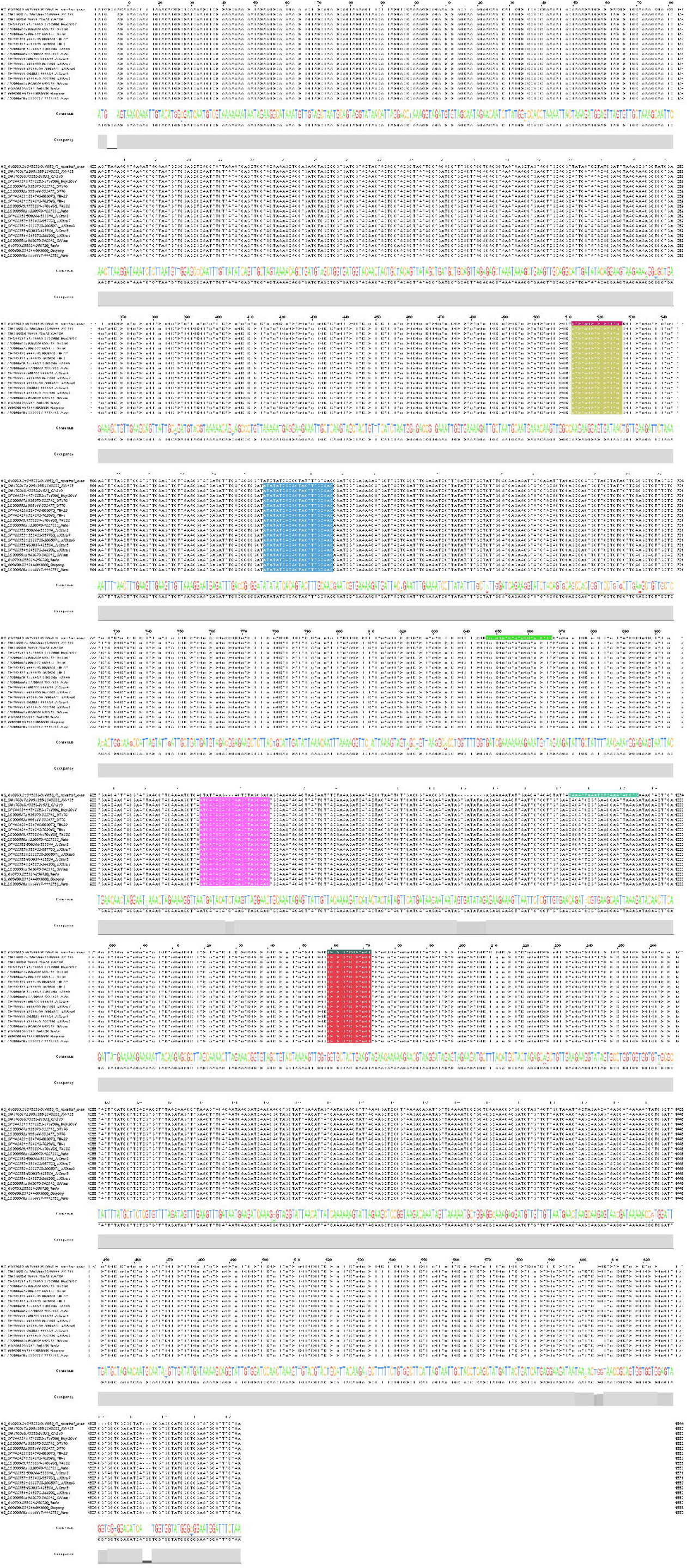
PCR primer binding site on groEL gene of different strains of *O. tsutsugamushi*. Strain information is provided in Table 4. The Gro-1 and Gro-2 primer-binding sites for *Rickettsia rickettsii* are shown in purple and dark green, respectively, whereas those for *O. tsutsugamushi* are shown in yellow and red, respectively. The SF1 and SR2 primer-binding sites are shown in different shades of green and bind only to *R. rickettsii* (NC_010263.3). The TF1 and TF2 primer-binding sites are shown in blue and rose, respectively, and bind only to *O. tsutsugamushi*.

## Conclusion

Taken together, our findings suggest that nested PCR targeting the *groEL* gene may be a useful and reliable approach for the detection and identification of *O. tsutsugamushi* in ectoparasite samples.

## Author contribution

**RG:** Formal analysis (Field collection, Identification); Writing – review & editing. **SGS:** Investigation; Methodology; Formal analysis; Writing – original draft, review & editing. **RM:** Writing – review & editing. **AAPA:** Conceptualization (Bioinformatics); Data analysis; Validation; Writing – original draft, review & editing. **PPS:** Supervision; Writing – review & editing. All authors reviewed and approved of the final manuscript.

## Acknowledgements

The authors thank the Director of ICMR-NIVCR for providing all the necessary amenities, guidance, encouragement, and useful suggestions that enabled the completion of this study. We would like to extend our sincere gratitude to all the ICMR-NIVCR Field Station staff in Madurai, Tamil Nadu, for their support in this study. AAPA acknowledges DBT-BIF Centre, Lady Doak College for providing bioinformatics facility.

## Ethics declaration

The study was approved by the Institutional Animal Ethics Committee of Madurai Medical College, Madurai, Tamil Nadu.

